# Membrane remodeling properties of the Parkinson’s disease protein LRRK2

**DOI:** 10.1101/2022.08.10.503505

**Authors:** Xinbo Wang, Javier Espadas, Yumei Wu, Shujun Cai, Jinghua Ge, Lin Shao, Aurélien Roux, Pietro De Camilli

**Affiliations:** Department of Neuroscience, Yale University School of Medicine, New Haven, Connecticut 06510, USA; Department of Cell Biology, Yale University School of Medicine, New Haven, Connecticut 06510, USA; Howard Hughes Medical Institute, Yale University School of Medicine, New Haven, Connecticut 06510, USA; Program in Cellular Neuroscience, Neurodegeneration and Repair, Yale University School of Medicine, New Haven, Connecticut 06510, USA; Kavli Institute for Neuroscience, Yale University School of Medicine, New Haven, Connecticut 06510, USA; Aligning Science Across Parkinson’s (ASAP) Collaborative Research Network, Chevy Chase, MD, 20815, USA; Department of Biochemistry, University of Geneva, Geneva, CH-1211, Switzerland

**Keywords:** LRRK2, Parkinson, tubulation, GTPase, membrane curvature

## Abstract

Mutations in Leucine-rich repeat kinase 2 (LRRK2) are responsible for late-onset autosomal dominant Parkinson’s disease (PD). LRRK2 has been implicated in a wide range of physiological processes including membrane repair in the endolysosomal system. Here, using cell free systems, we report that purified LRRK2 directly binds acidic lipid bilayers with a preference for highly curved bilayers. While this binding is nucleotide independent, LRRK2 can also deform low curvature liposomes into narrow tubules in a guanylnucleotide-dependent but ATP-independent way. Moreover, assembly of LRRK2 into scaffolds at the surface of lipid tubules can constrict them. We suggest that an interplay between the membrane remodeling and signaling properties of LRRK2 may be key to its physiological function. LRRK2, via its kinase activity, may achieve its signaling role at sites where membrane remodeling occurs.

**Significance Statement:** LRRK2 is one of the most frequently mutated gene in familial Parkinson’s disease. While much has been learned about its molecular properties, upstream regulators and protein substrates of its kinase activity, its precise function remains unclear. Recent evidence has pointed to a role of LRRK2 in membrane repair in the endo/lysosomal system. Here we show that purified LRRK2 has membrane remodeling properties. We suggest that its ability to sense and induce membrane curvature may be key to its function in membrane dynamics. These properties may help coordinate a direct role of LRRK2 at the membrane interface with its the signaling role of its kinase domain.

## Introduction

LRRK2 is the most frequently mutated gene in familial Parkinson’s disease (PD) and mutations in LRRK2 are associated with increased risk for the disease (1–4). It is a multi-domain protein that contains two catalytic modules, a Roc (Ras of complex) GTPase domain and a serine-threonine kinase domain separated by a COR (C-terminal of Roc) domain. These two domains are flanked by four protein-protein interaction modules: Armadillo, Ankyrin, Leucine-Rich Repeats at the N-terminus and a WD40 domain at the C-terminus (Fig. 1A) (5–7). Growing evidence suggests a role of LRRK2 in membrane dynamics in the endolysosomal system mediated at least in part by its property to phosphorylate Rab proteins (8–14). During lysosome stress, LRRK2 is recruited to damaged lysosomes where it has been proposed to function in membrane repair and membrane traffic reactions (10, 15–17). A variety of other roles for LRRK2 have also been proposed (9, 18).

**Fig. 1.**
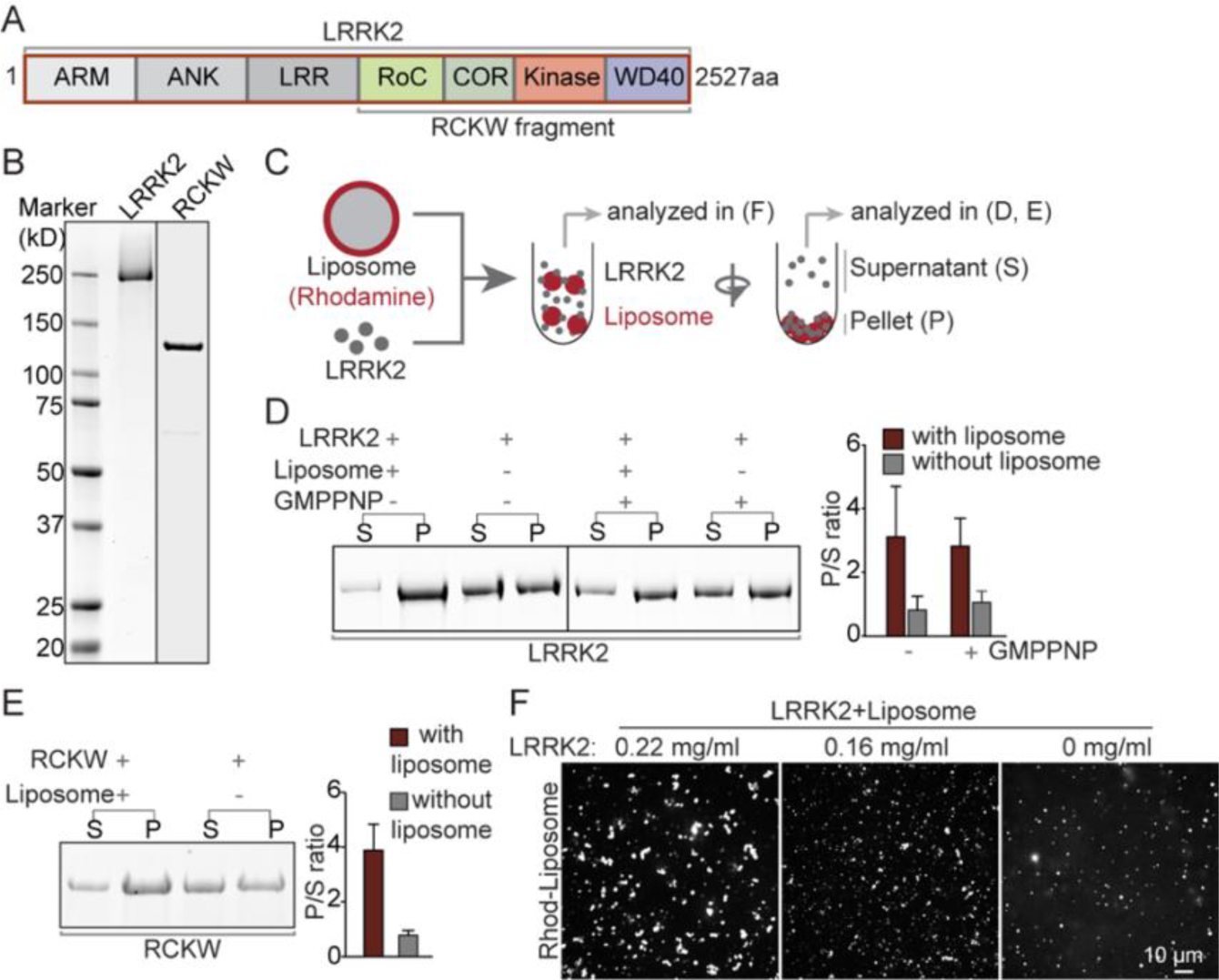
Purified LRRK2 binds liposomes. (A) Domain cartoon of full length human LRRK2. (B) LRRK2 and the RCKW fragment were purified from Expi293 cells and analyzed by SDS-PAGE and Coomassie Blue (CB) staining. (C) Schematic diagram of the protein-liposome binding assay. (D and E) Left panels, Supernatants (S) and pellets (P) of the centrifugation step were analyzed by SDS-PGE and Coomassie Blue staining. Right panel, Quantification of the intensities of the gel bands as shown in the left panels. Bars represent the ratio between pellets and supernatants (P/S). Values represent mean ± S.D. n≥3 independent experiments. (F) Confocal microscopy analysis of rhodamine (Rhod)-labeled liposomes incubated with different concentrations of LRRK2, showing LRRK2-induced liposome clusters.

When overexpressed in cultured cells, some PD-causing LRRK2 mutant protein polymerize into helices around microtubules (5, 19). Moreover, the purified RCKW C-terminal fragment of LRRK2 (Fig. 1A) binds and assembles into helices around microtubules *in vitro* (6, 20). These properties have facilitated the characterization of the atomic structure of LRRK2 by cryo-EM (6, 20) and cryo-electron tomography (cryo-ET) (5), but the physiological significance of LRRK2 assembly on microtubules remains unclear. The microtubule binding properties of LRRK2 share similarities with dynamin, another helix-forming GTPase, which had originally been shown to assemble into helices around microtubules (21, 22). However, subsequent research revealed that dynamin has membrane remodeling properties and that the physiological templates for dynamin assembly are endocytic membrane tubules (23–26). In view of evidence linking LRRK2 to events occurring at membrane surfaces and the precedent set by dynamin, we hypothesized that the negatively charged surface of microtubules may mimic a negatively charged tubular membrane template. Thus, although LRRK2 is structurally different from dynamin, we set out to determine whether LRRK2 can bind and remodel lipid membranes and assemble around lipid tubules.

## Results

### LRRK2 binds lipid bilayers *in vitro*

Full-length flag-tagged LRRK2 or its flag-tagged C-terminal RCKW fragment were expressed in Expi293 cells, purified by anti-flag affinity-purification and then used after cleavage of the tag (Fig. 1B and Fig. S1A-C). Purified LRRK2 was enzymatically functional as it had both GTPase activity (Fig. S1D) [a lower activity than dynamin, but in line with data from the literature (27, 28)] and kinase activity (Fig. S1E), as determined by its property to phosphorylate itself and Rab8, one of its known substrates (8). To assess LRRK2’s membrane binding ability, either LRRK2 (300 nM) or the RCKW fragment (300 nM) were incubated at 37^0^C for 30 minutes in physiological salt and pH together with acidic liposomes (20 μM) and then subjected to centrifugation (Fig. 1C). As shown by SDS-PAGE analysis, the presence of liposomes strongly enhanced the fraction of LRRK2 and RCKW recovered in the pellet (Fig. 1D, E) revealing that LRRK2 binds acidic lipid membranes. In separate experiments, the incubation of LRRK2 with rhodamine (Rhod)-labeled liposomes was carried out on coverslips under microscopic observations. Such analysis revealed that LRRK2 clustered liposomes in a concentration-dependent way (Fig. 1F), further supporting its binding to membrane.

### LRRK2 deforms liposome into tubules

We next tested the effect of nucleotides on LRRK2 binding to liposomes as its GTPase and kinase domains are critical to its physiological function (29). Presence of GMPPNP, a nonhydrolyzable GTP analog, did not affect the co-sedimentation of LRRK2 with liposomes (Fig. 1D). However, microscopic examination revealed that, in the presence of GMPPNP, LRRK2 promoted the robust formation of membrane tubules from liposomes (Fig. 2A). Thus, GMPPNP, while lacking an effect on the liposome binding properties of LRRK2, unmasks membrane remodeling properties of this protein. Tubulation was “all or none”, with the formation of very long tubules from a small subset of large liposomes or liposome clusters, suggesting a nucleation-dependent process. Similar results were also obtained with the RCKW fragment (Fig. 2B), indicating that all the protein determinants required for membrane deformation are contained in this fragment, as in the case of binding to microtubules (5, 6, 20). Replacing unlabeled proteins with GFP tagged-LRRK2 or Alexa 488 labeled-RCKW demonstrated precise colocalization of the fluorescent proteins with the labeled lipids (Fig. 2C, D and Fig. S2A, B), confirming that the lipid tubules were indeed coated by either LRRK2 or RCKW. Moreover, dynamic imaging of GFP tagged-LRRK2 and Rhod-labeled liposomes showed that tubule elongation correlated with LRRK2 presence on their surface (Fig. 2E and movie S1). Upon photobleaching, Rhod-liposome fluorescence recovered quickly within seconds, whereas fluorescence of GFP-LRRK2 did not (Fig. S2C), indicating the formation of stable assemblies on the tubular membrane.

**Fig. 2.**
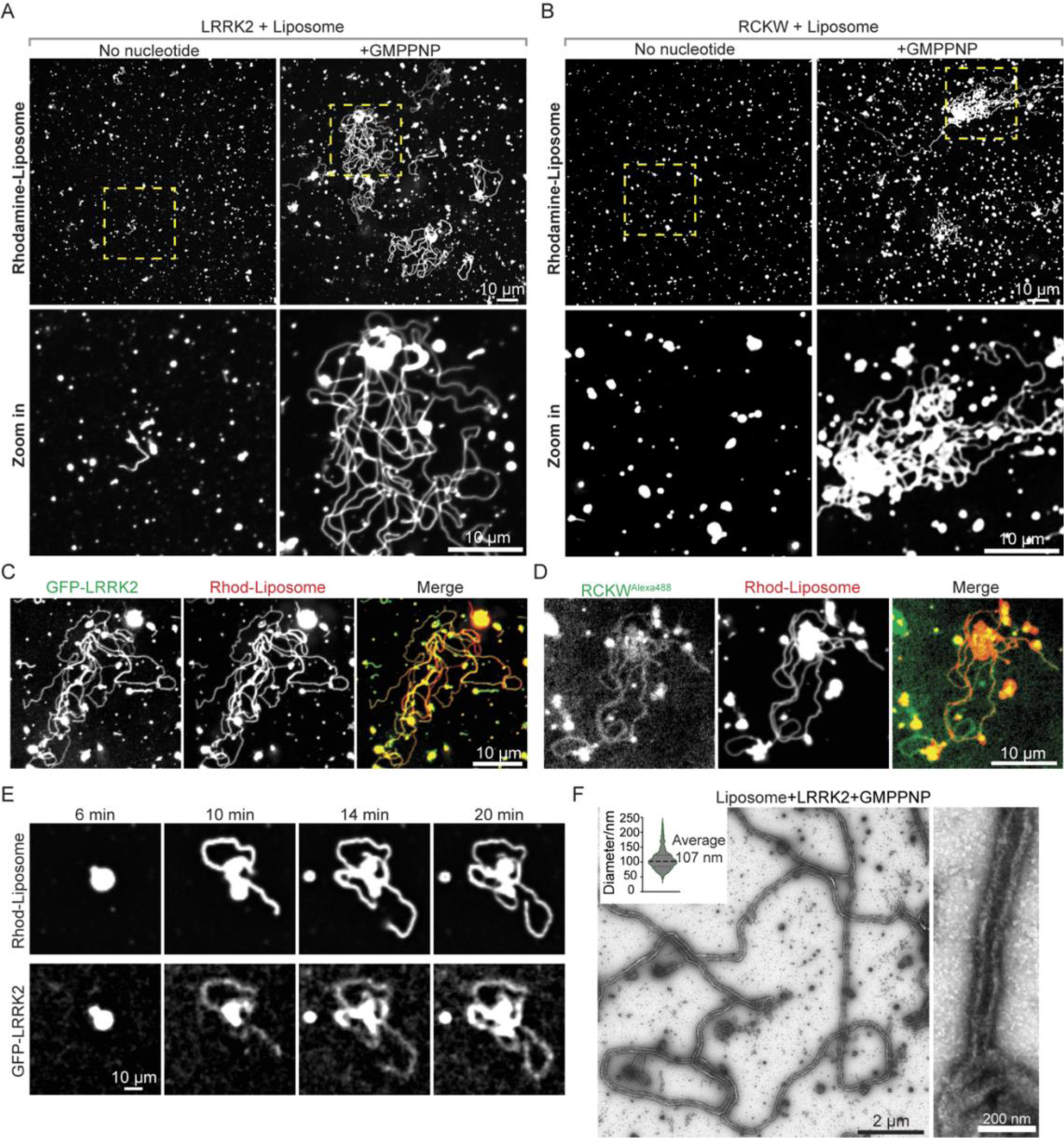
LRRK2 has membrane remodeling properties. (A and B) Confocal microscopy analysis of Rhodamine-labeled liposomes incubated with purified full length LRRK2 (A) or the RCKW fragment (B) in the presence (right panels) or absence (left panel) of GMPPNP for 30 mins at 37^0^C. Lower panels show the regions indicated by boxes in the upper panel at higher magnification. (C and D) Fluorescence images of Rhod-labeled liposomes incubated with full length GFP-LRRK2 (C) or Alexa488-RCKW (D) in the presence of GMPPNP. (E) Representative time-lapse sequence showing emergence and elongation of Rhod-labeled liposome tubules coated by GFP-LRRK2. (F) EM micrographs showing negative stained liposome tubules induced by LRRK2 in the presence of GMPPNP. The inset at top left shows quantification of diameters of LRRK2-coated tubules (diameters measured include the protein coat). One high magnification image is shown on the right.

Analysis of mixtures of full length LRRK2 with liposomes in the presence of GMPPNP by negative staining EM confirmed the presence of narrow lipid tubules with somewhat variable diameter (average outer diameter of the coated tubule = 107 nm) (Fig. 2F).

An algorithm to quantify liposome tubulation in fluorescence images was developed to quantify the effect of various nucleotides on this process (see methods) (Fig. 3A). This quantification validated the effect of GMPPNP on tubulation (Fig. 3B, C) and also revealed that this property was shared by other guanine nucleotides tested, but not by ATP (Fig. 3B and Fig. S2D). GTP had similar efficiency to its non-hydrolyzable analogues GMPPNP and GTPγS, and GDP was as efficient as GTP (Fig. 3B and Fig. S2D), demonstrating that guanine nucleotides binding, but not hydrolysis, is important for the membrane remodeling properties of LRRK2 in our *in vitro* system.

**Fig. 3.**
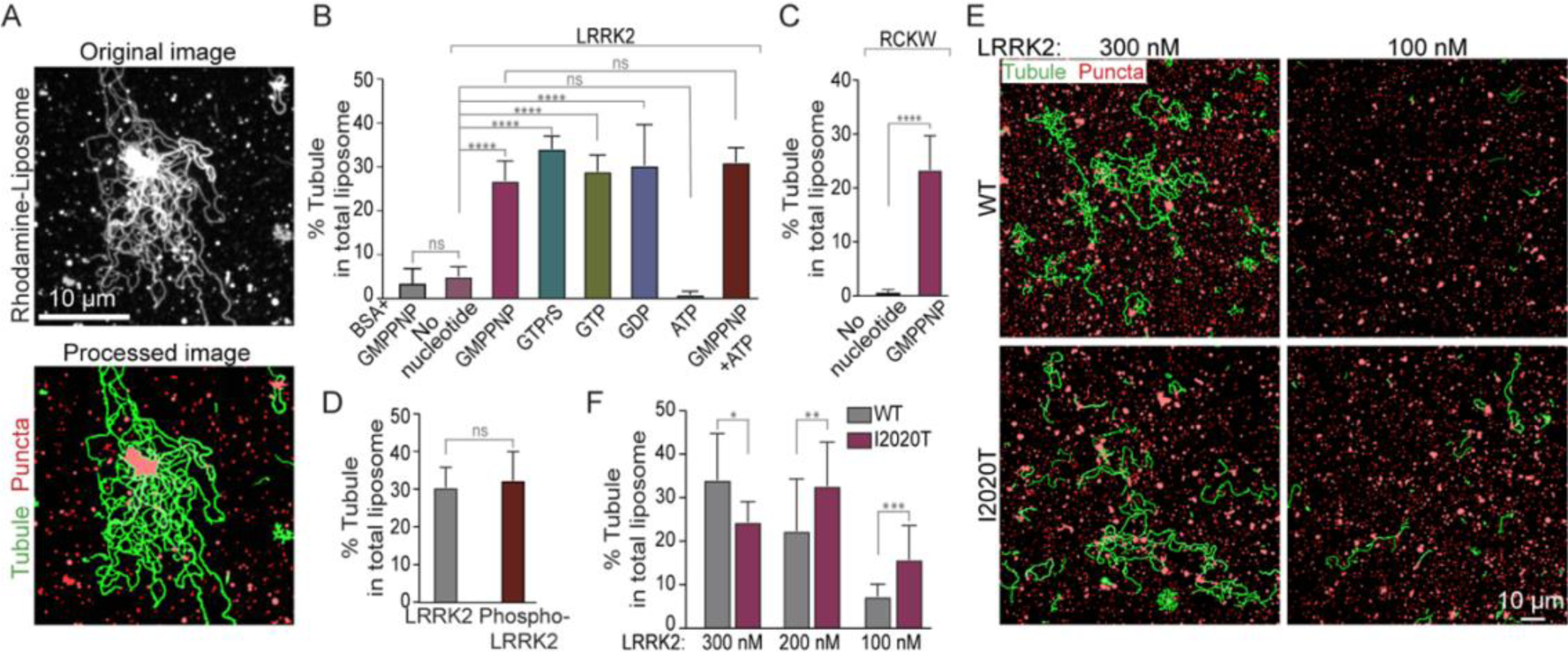
Conditions that impact LRRK2-dependent liposome tubulation. (A) A software was generated to enhance visualization of tubules in order to allow their quantification. Representative fluorescence micrographs showing LRRK2-induced liposome tubulation in the presence of GMPPNP before (top) and after image processing (bottom). Liposome tubules are labeled in green, non-tubular liposomes are labeled in red. (B and C) Quantification of the tubulation efficiency of full length LRRK2 (B) or the RCKW fragment (C) in the presence of different nucleotides. (D) Quantification of the tubulation efficiency of autophosphorylated LRRK2, showing that autophosphorylation does not affect tubulation efficiency. (E) Representative fields, after image processing to highlight membrane tubules, of liposome incubated with wild-type (upper) and kinase domain mutant (I2020T) LRRK2, at the concentration indicated. (F) Quantification of the tubulation efficiency of WT and mutant LRRK2^I2020T^ at the indicated protein concentration. Bars represent mean ± S.D. n≥10 images from at least 3 independent experiments. Data from (B) and (F) were analyzed using Ordinary one-way ANOVA by Prism 9; Data from (C) and (D) were analyzed using t-tests by Prism 9. ****, P < 0.0001; ***, P < 0.001; **, P < 0.01; *, P < 0.05; ns, P > 0.05

The kinase activity of LRRK2 is critical for its physiological function. Many of the LRRK2 mutations that cause, or increase the risk of PD increase LRRK2 kinase activity (9, 29, 30). However, as shown above, ATP had no effect on LRRK2-dependent tubulation (Fig. 3B), indicating that ATP binding and kinase activity of LRRK2, leading to its phosphorylation, do not contribute to its property to deform liposomes *in vitro*. Additionally, autophosphorylated LRRK2 that had been generated by an *in vitro* phosphorylation reaction, and whose autophosphorylated state was confirmed by western blotting (Fig. S3A), was not more potent than LRRK2 not subjected to this reaction in inducing liposome tubulation (Fig. 3D and Fig. S3B).

The PD-causing mutation I2020T in the kinase domain of LRRK2 (LRRK2^I2020T^) enhances its protein kinase activity and its polymerization around microtubules (5, 6, 19, 20). The same mutation also increased the liposome tubulation property of LRRK2 in our system which does not include ATP (Fig. 3E, F and Fig. S3C), without affecting its membrane binding (Fig. S3D). This enhancement was stronger at lower LRRK2 concentrations (Fig. 3E, F), probably reflecting the lower critical concentration required for the nucleation of LRRK2 assemblies on the membrane. Considering that autophosphorylation of LRRK2 does not affect its membrane remodeling properties, the I2020T mutation may act by an effect on the conformation of the kinase domain which in turn affects the conformation of the entire protein, as previously suggested to explain enhanced microtubule binding by this mutant (19, 31). It was proposed that the N-terminal moiety of LRRK2, comprising the ARM, ANK, and LRR domains (Fig. 1A), has an inhibitory regulatory role on its kinase module, by acting as a lid that locks it into an autoinhibited state (31). Releasing this autoinhibitory state is an essential prerequisite for microtubule binding and polymerization.

### LRRK2 assembles around preformed narrow lipid tubules

To test more directly the possibility that binding of LRRK2 to microtubules may mimic binding to tubular membrane templates, we explored binding of LRRK2 to preformed highly curved bilayers with a width in the same range as microtubules. Narrow lipid-nanotubes were prepared as described (32) by using a lipid mixture containing in moles 40% galactosylceramide (GC) and 60% PtdSer (PS) (GC/PS liposomes). Negative stain EM showed that this mixture preferentially assembled into lipid-nanotubes with fixed geometry and diameter (outer diameter of the bilayer ∼ 29 nm) similar to diameter of microtubules (Fig. S4B), although some non-tubular liposomes were also observed (Fig. S4B). Negative staining EM (Fig. 4A) of GC/PS nanotubes (20 μM) incubated with LRRK2 (300 nM) demonstrated presence of a dense continuous coat on their surface (Fig. 4A), irrespective of the presence of nucleotides. A regular pattern of the coat could not be observed, but the presence of naked regions next to coated regions (Fig. 4A) was consistent with the known property of LRRK2 to polymerize (5, 6, 20). No such coat was observed on nanotubes not incubated with LRRK2 (Fig. 4A). As reported for LRRK2 coated microtubules (5), LRRK2 coated lipid-nanotubes had the propensity to bundle (Fig. 4A and Fig. S4A). Coating of lipid nanotubules by LRRK2 was also observed by cryo-EM, using for the incubation with LRRK2 conditions that closely reflected those used to characterize binding of LRRK2 to microtubules (Fig. 4B). The property of LRRK2 to assemble on GC/PS nanotubes was confirmed by fluorescence experiments involving incubation of GFP-LRRK2 (100 nM) with these nanotubes labeled with trace amounts of Cy5-labelled-PE. These experiments additionally revealed a preferential binding of LRRK2 to the nanotubes relative to non-tubular liposomes present in the sample (Fig. 4C), indicating that LRRK2 prefers to bind high curvature membranes.

**Fig. 4.**
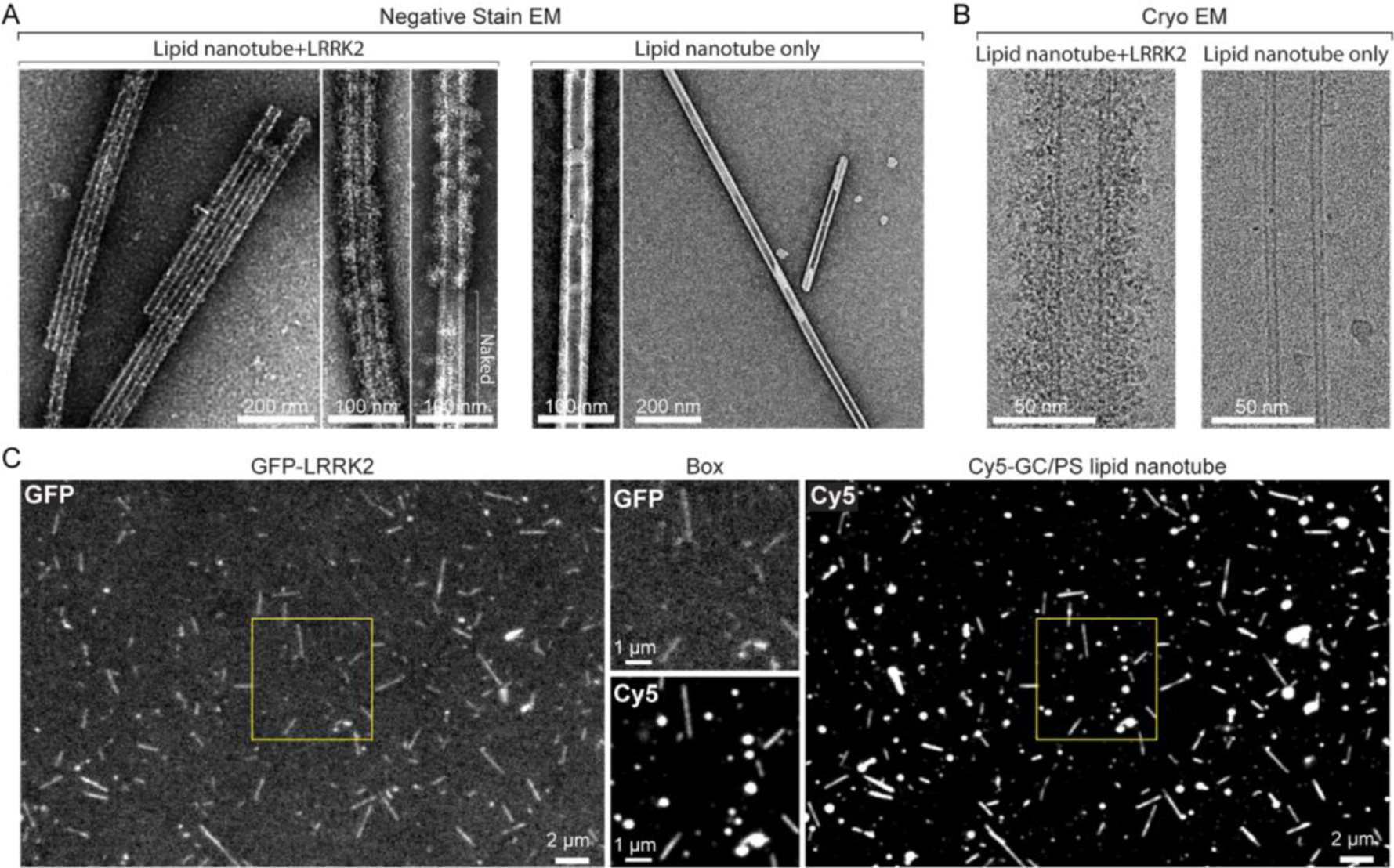
Assembly of LRRK2 around preformed lipid nanotubes. (A) EM images of negatively stained lipid nanotubes (GC/PS liposomes) incubated with LRRK2 (left), and of lipid nanotubes only as a control (right). High magnification images are also shown. A dense coat is present on nanotubes incubated with LRRK2 (but note a naked portion of the nanotube in one of the high magnification images). The low magnification image at left shows that presence of LRRK2 also induce nanotube bundling. (B) Cryo-EM images of LRRK2 assembled on a lipid nanotube (left) and of a nanotube only control (right). (C) Fluorescence images of GC/PS liposomes labeled with Cy5 incubated with GFP-LRRK2. The regions boxed by yellow squares are shown at higher magnification in the two middle images. Comparison of the Cy5 and GFP fluorescence reveals preferential binding of LRRK2 to lipid nanotubes.

To further determine whether LRRK2 has membrane curvature sensing properties, we analyzed its binding to narrow tubules pulled from giant unilamellar vesicles (GUVs) composed in moles by 59.9% DOPC, 40% DOPS and 0.1% Atto647N DOPE. In this system, the diameter of the tubules could be calculated by generating a calibration curve that relates the fluorescence of Atto647N DOPE to tubule diameter. Light microscopy analysis of these tubules exposed to low concentrations of GFP-LRRK2 (20 nM) showed preferential binding of LRRK2 to the tubules relative to the shallow surface of the GUVs (Fig. 5A). The curvature preference was quantified by determining a sorting ratio defined as the ratio between the protein density (GFP-LRRK2 fluorescence relative to the fluorescence of Atto 647N DOPE) on the tubules and the protein density on the surface of the GUV (see Methods). This sorting ratio revealed a preference of LRRK2 for tubules and more so for narrow tubules compared to wider ones (Fig. 5B), with no significant binding difference being observed based on the absence or presence of nucleotides (ATP, GTP or GDP) (Fig. 5A, B).

**Fig. 5.**
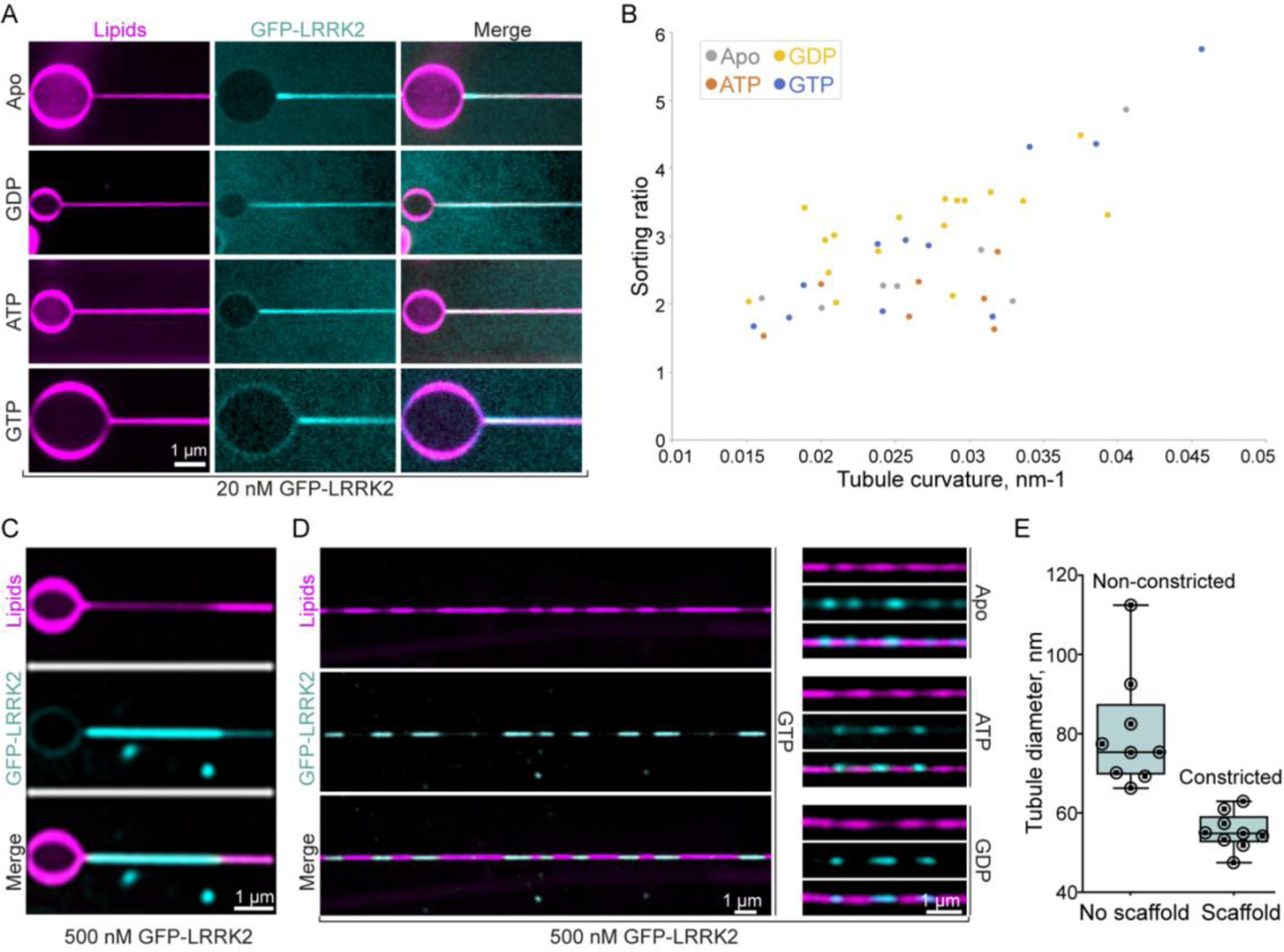
LRRK2 has membrane curvature sensing and generating properties. (A) and (B) Narrow lipid tubules were pulled from GUVs composed of an acid phospholipid mixture and exposed to a low concentration (20 nM) of GFP-LRRK2 in the presence or absence of the indicated nucleotides (5 mM). (A) Representative fluorescence images demonstrating that GFP-LRRK2 is preferentially recruited to the tubules revealing that it has curvature sensing properties. (B) Plot showing the sorting ratio of GFP-LRRK2 on individual tubules. Ratios were defined as the ratios between the protein density (GFP-LRRK2 fluorescence relative to the fluorescence of Atto 647N DOPE) on the tubules and the protein density on the surface of the GUV. A calibration curve allowed to derive the tubule diameter from the lipid fluorescence. Each data point corresponds to a different membrane tubule. (C) - (E) When used at high concentration (500 nm), GFP-LRRK2 assembles into high density scaffolds on the surface of the tubules, leading to membrane constriction. (C) Representative example of a GFP-LRRK2 scaffold formed on a lipid tubule pulled from a GUV, in the presence of GTP (5 mM). (D) Examples of GFP-LRRK2 scaffolds on lipid tubules generated by the lipid covered silica bead system, in the presence or absence of the indicated nucleotides (5 mM). (E) Quantification of tubule diameter in segments covered or non-covered by the GFP-LRRK2 scaffolds, in the presence of GTP (5 mM).

Importantly, upon exposure to high GFP-LRRK2 concentration (500 nM), tubule-bound LRRK2 segregated itself into regions where it was present at much higher density than in surrounding portions of the tubule. In these regions there was a corresponding decrease of the lipid fluorescence (Atto 647N DOPE) indicating formation of a LRRK2 scaffold that constricts the underlying tubule (Fig. 5C).

To further analyze these LRRK2 assemblies on lipid tubules in a regime of high LRRK2 concentration, we employed an experimental system in which long narrow lipid tubules of variable diameters are generated by lipid-covered silica beads rolling on an inclined glass (see Methods). The same lipid composition used for tubules pulled from GUVs was used in this system. When these tubules were generated in the presence of GFP-LRRK2 (500nM), discrete segments of intense GFP-LRRK2 fluorescence were observed along the tubules, irrespective of the presence of nucleotides (Fig. 5D), although in the presence of GTP this effect was more pronounced and consistent. In these segments reduced lipid fluorescence was observed, suggesting that presence of LRRK2 scaffolds had constricted the tubules (Fig. 5D). Based on the calibration curve with Atto 647N DOPE fluorescence, the diameters of lipid tubules in the regions of low GFP-LRRK2 fluorescence were substantially variable (range: from 66.4 nm to 112.6 nm; mean 80.2 nm). In contrast, in the segments of intense GFP-LRRK2 fluorescence, tubules were thinner and of quite uniform diameter (range: from 52.0 nm to 63.0 nm: mean 55.4 nm) confirming that, as in the case of the GUV-based assay, the assembly of LRRK2 scaffolds had produced membrane constriction. Moreover, the homogenous diameters of the constricted tubules suggest that LRRK2 assembles into a scaffold with a specific geometry (Fig. 5E). The assembly of LRRK2 scaffolds that constrict lipid tubules is reminiscent of what has been observed with dynamin with similar experimental systems (33, 34), although scaffold formation in the case of dynamin is followed by fission upon addition of GTP (33, 34).

## Discussion

In this study we investigated the interaction between LRRK2 and lipid bilayers using four different *in vitro* membrane systems: spherical liposomes, preformed lipid nanotubes, lipid tubules pulled from GUVs, and lipid tubules generated by membrane coated silica beads rolling. We demonstrate that LRRK2 can directly bind to lipid bilayers regardless of its nucleotide binding state, that it shows preference for high curvature membranes and that it can also constrict preformed membrane tubules. We further observed that LRRK2 can deform low curvature liposomes into narrow membrane tubules, but that this property requires LRRK2 to be in its guanylnucleotide-bound state, revealing that a pre-existing high membrane curvature can override guanylnucleotide requirement for LRRK2 assembly. Guanylnucleotide binding may help the protein acquire a conformation that reduces its resistance to polymerization into a tubular structure after its binding to low curvature membranes. A role of high membrane curvature in facilitating membrane binding was also observed for dynamin (33, 34).

The tubules generated by LRRK2 from low curvature liposomes have variable diameters. Moreover, the diameter of the preformed lipid nanotubes is smaller than the average diameter of tubules induced by LRRK2 from liposomes. These results suggest that LRRK2 has conformational flexibility, enabling it to accommodate different membrane geometries. This flexibility is also evident in its binding to microtubules, as LRRK2 can assemble into a helical coat on microtubules with different number of protofilaments (5). However, when added to preformed tubules at high concentration, LRRK2 appears to assemble into a scaffold with a specific geometry, as indicated by the relatively homogeneous lipid tubule diameter beneath the scaffolds (Fig. 5D, E). While it would be interesting to examine the structure of this LRRK2 scaffold by Cryo-EM, technical limitations prevented us so far from generating these tubules on EM grids for this analysis.

Collectively, our results show that, as observed for several proteins implicated in membrane remodeling (35–37), LRRK2 has curvature sensing and curvature inducing properties. In living cells, direct binding of LRRK2 to membranes may cooperate with other factors, such as Rab proteins, which may provide spatial specificity in its membrane recruitment (13, 38, 39). A membrane remodeling property of LRRK2 is consistent with studies implicating LRRK2 in membrane repair in the lysosomal system (10, 15, 16) and with its functional partnership with proteins, such as Rabs (8, 39), which function at sites of membrane dynamics. While a functional significance of the reported binding of LRRK2 to microtubules cannot be excluded – f.e. PD-causing mutations that increase affinity for microtubules could interference with microtubule-dependent transport (6) -, most current evidence favors a function of LRRK2 at membrane interfaces. Most interestingly, in connection to our findings, LRRK2 was reported to trigger the formation of JIP4-positive membrane tubules from lysosomes upon rupture of their membrane by the lysosomotropic reagent LLOMe (L-Leucyl-L-leucine methyl ester) (10, 40), although LRRK2 itself was not detected on such tubules. LRRK2 may help nucleate a membrane tubule that is then elongated by other factors. Other membrane remodeling proteins that can induce long narrow tubules from liposomes *in vitro*, function in living cells only by forming transient assemblies on very short tubular intermediates not recognizable as tubules by light microscopy (35, 37, 41).

In conclusion, our findings raise the possibility that membrane remodeling properties of LRRK2 assemblies may play a role in its function. Such properties may cooperate with the signaling properties of LRRK2 and may help control where such signaling occurs.

## Materials and Methods

### Materials, Reagents and Antibodies

ExpiFectamine293 Transfection Kit, Thermo Scientific, A14525; Prescission Protease, GenScript, Z02799; 3xFLAG Peptide, Sigma, F4799; Protease inhibitor cocktail, Roche, 05056489001; Glutathione Sepharose, GE Healthcare, 17075601; Mini dialysis units, Thermo Scientific, 69572; Centrifugal filters units, Sigma, UFC901024; SuperSep Phos-tag Gel, FUJIFILM, 195-1799; 35mm glass bottom dishes, MatTek, P35G-1.5-14-C; Alexa Fluor 488 Protein Labeling Kit, Thermo Scientific, A10235; PD-10 Desalting Columns, GE Healthcare, 17085101; EnzChek Phosphate Assay Kit, Invitrogen, E6646; His60 Ni Superflow Resin, Takara Bio, 635660; MLi-2, Tocris, 5756; Isopropyl b-D-thiogalactoside (IPTG), AmericanBio, 367-93-1; Carbon film mesh, Electron Microscopy Sciences, 2154128400. All lipids were obtained from Avanti Polar Lipids: Brain PS, 840032; Rhod-PE, 810150; Cy5-PE, 810345; Galactosylceramide, 860546P. All nucleotides were obtained from Sigma-Aldrich: Guanosine 5’-[beta, gamma-imido]triphosphate trisodium salt hydrate (GMP-PNP), G0635; Guanosine 5’-triphosphate sodium salt hydrate (GTP), G8877; Guanosine 5’-diphosphate sodium salt (GDP), G7127; GTP-gamma-S,Tetralithium salt (GTPrS), 10220647001 ; Adenosine 5′-triphosphate disodium salt hydrate (ATP), A2383. Anti-FLAG M2 Affinity Gel, Sigma, A2220; Anti-LRRK2 (phospho T1357) antibody, Abcam, ab270606; Monoclonal ANTI-FLAG M2 antibody, Sigma, F3165; Rab8A Rabbit mAb, Cell signaling, 6975.

### Plasmids

Human full length wild-type LRRK2 (LRRK2-FL) (NM_198578.4) within the p3xFLAG-CMV10 vector was a kind gift from Karin Reinisch (Yale University, New Haven, CT). This construct was further modified by inserting a PreScission Protease recognition sequence between the N-terminal 3xflag tag and the LRRK2 sequence through enzyme digestion and ligation. The LRRK2-I2020T mutation was generated through site-directed mutagenesis of this construct by PCR. The sequence encoding the C-terminal fragment (RCKW) of LRRK2 was obtained by PCR from the same construct and then inserted into the p3xFLAG-CMV10 vector through enzyme digestions and ligation between the NotI/BamHI restriction sites. For the GFP-LRRK2 fusion construct, the sequence coding for the enhanced green fluorescent protein (eGFP) was fused in frame to the N-terminus of LRRK2 between the PreScission recognition sequence and NotI restriction site using HiFi assembly (NEB). A plasmid encoding Rab8 was previously generated in the De Camilli lab.

### Protein expression and purification

LRRK2. Constructs encoding 3xFlag-LRRK2, 3xFlag-LRRK2(I2020T), 3xFlag-RCKW or 3xFlag-GFP-LRRK2 were transfected into Expi293F cells (Thermo Fisher Scientific) according to manufacturer instructions. Proteins were expressed for three days following induction. Cells were harvested by centrifugation and used immediately for protein purification. Cells were lysed by 3 freeze-thaw cycles in lysis buffer that contained 20mM HEPES 7.4, 500 mM NaCl, 10% Glycerol, 2mM DTT and 1xcomplete EDTA-free protease inhibitor cocktail (Roche). Cellular debris were removed by centrifugation at 15000xg for 1 hour at 4°C, and the clarified lysate was mixed with anti-FLAG M2 resin (Sigma) for 2 h while rotating at 4°C. The resin was then washed with 3x10 bed volumes of lysis buffer and eluted with 800 µL (for 60 ml of cell suspension) lysis buffer (without protease inhibitor) supplemented with 0.2 mg/mL 3x Flag peptides (Sigma). The N-terminal 3xFlag tag was removed by incubation with the GST tagged Prescission Protease (GenScript) (final 0.01U/ul) overnight at 4°C and the GST tagged Prescission Protease was subsequently removed by Glutathione Sepharose. The purity of the proteins was assessed by SDS-PAGE and Western blotting. Purified proteins were dialyzed overnight at 4°C against a buffer containing 20mM HEPES 7.4, 150 mM NaCl, 2.5 mM MgCl2, 5% Glycerol, 2mMDTT, 20 μM GDP. After dialysis proteins were further clarified by centrifugation at 17000xg for 10 min at 4°C, their concentration was determined by SDS-PAGE using Bovine Serum Albumin (BSA) as standard and used without freezing in liposome binding and tubulation experiments. The detailed protocol was deposited in protocols.io (DOI: dx.doi.org/10.17504/protocols.io.8epv59wd4g1b/v1)

Dynamin. Dynamin was purified from rat brain extract as described (42). Brain extract was prepared by homogenizing rat brains in a glass-Teflon homogenizer in lysis buffer (10 ml/brain) that contained 20 mM HEPES pH 7.4, 150 mM NaCl, 4 mM DTT, protease inhibitor cocktail (Roche). Triton X-100 was then added (final concentration 0.1%). Insoluble material was removed by centrifugation at 15 000xg for 1 hour at 4°C. The clarified brain extract was incubated with the GST-tagged SH3 domain of rat amphiphysin 2 as an affinity ligand as described (43). After washing extensively with the lysis buffer, dynamin was eluted with elution buffer that contained 20 mM PIPES pH 6.2, 1.2 M NaCl, 10 mM Ca2+, 1 mM DTT, and dialyzed overnight against dialysis buffer (20 mM HEPES, 100 mM NaCl, 50% Glycerol, 2mM DTT). Purity of the protein was assessed by SDS-PAGE and Coomassie Blue staining.

Rab8. Rab8 was purified from bacteria as described (8). Constructs encoding 6xHis-Rab8 were transformed into M15 Competent Cells. Cells were grown in Super Broth medium at 37°C to an OD600 of 0.6, and Rab8 expression was induced by addition of 0.5 mM IPTG for 18 hours at 18°C. Cells were harvested and lysed by sonication in lysis buffer that contained 50 mM phosphate pH 7.5, 150 mM NaCl, 10% glycerol, 5 mM MgCl2. Lysates were further clarified by centrifugation at 15000 × g for 1 h, the protein was isolated by a Ni-NTA column, and further purified by gel filtration in storage buffer (10 mM HEPES pH 7.5, 150 mM NaCl, 5 mM MgCl2 and 2 mM DTT).

### Protein labeling

3xflag-tagged RCKW was purified as described above. Conjugation of RCKW to Alexa 488 was carried out according to the Alexa Fluor 488 Protein Labeling Kit [Thermo Scientific, using conditions aimed at achieving low degree of labeling (DOL<25), see the manufacturer instructions for the details], with the exception that after cleavage of the flag tag in lysis buffer, the GST-Prescission Protease was removed by Glutathione Sepharose as described above. Free Flag peptides were removed using a PD-10 Desalting Column (GE Healthcare). The conjugate was further purified by a new PD-10 Desalting Column, concentrated by centrifugal filters (Sigma), dialyzed into dialysis buffer and subsequently used without freezing.

### *In vitro* GTPase activity

GTPase assays were performed using the Enzchek phosphate assay kit (Invitrogen). Reactions were performed in a 100-μl volume with 5 μl 20× reaction buffer (1 M Tris-HCl, 20 mM MgCl2, pH7.5 and 2 mM sodium azide), 200 μM 2-amino-6-mercapto-7-methylpurine riboside, 0.1 U purine nucleoside phosphorylase, and 9 μM LRRK2 protein or 0.8 μM Dynamin 1. Samples were incubated for 30 min at 37°C in a 96-well plate (Corning). Reactions were initiated by the addition of 0.5 mM GTP. The absorbance at 360 nm was measured every 1 min over 45 min at 37°C by using a microplate reader (Synergy H1; BioTek). The detailed protocol was deposited in protocols.io (DOI: dx.doi.org/10.17504/protocols.io.q26g74qwqgwz/v1)

### *In vitro* kinase activity

LRRK2 kinase assays were performed with a final volume of 40 μL with 8 μg recombinant Rab8 and 200 nM LRRK2 proteins in kinase buffer that contained 20 mM Tris–HCl (pH7.5), 7.5 mM MgCl2 and 0.1 mM EGTA with or without 1 mM ATP. Assays were incubated for 2 hours at 30°C, quenched through addition of SDS sample loading buffer and heated at 95°C for 10 min. Reaction mixtures were resolved by SDS/PAGE or Phos-tag SDS/PAGE (FUJIFILM). Proteins were detected by Coomassie Blue staining or Western blot using antibodies of Rab8 and LRRK2, respectively. The detailed protocol was deposited in protocols.io (DOI: dx.doi.org/10.17504/protocols.io.kxygxzr2kv8j/v1)

### *In vitro* LRRK2 autophosphorylation

Autophosphorylation of LRRK2 was performed in a 1.5 mL Eppendorf tube with 1.4 mL purified LRRK2 protein obtained by elution from the anti-FLAG M2 resin as described in the protein purification section. The protein containing solution was mixed with 10X kinase buffer (same as above for the *in vitro* kinase assay) supplemented with 1mM ATP and GST-Prescission Protease (to remove the Flag tag) and incubated at 4°C overnight. GST-Prescission Protease was removed by Glutathione beads from the LRRK2 containing solution which was then concentrated by centrifugal filters (Sigma) and dialyzed overnight at 4°C against dialysis buffer (20mM HEPES 7.4, 150 mM NaCl, 2.5 mM MgCl2, 5% Glycerol, 2mMDTT, 20μM GDP). Autophosphorylation was further assessed by Western blotting using a LRRK2 phospho-specific (pT1357) antibody. Concentration of the protein was determined by Coomassie-stained SDS gel using Bovine Serum Albumin (BSA) as standard. The detailed protocol was deposited in protocols.io (DOI: dx.doi.org/10.17504/protocols.io.81wgb6m91lpk/v1)

### Liposome preparation

The composition of the liposome mixtures in moles percent was as follows: PS liposomes: 99.5% brain PS:0.5% Rhod-PE (Avanti, 810150).

GC/PS nanotubes: 39.5% Galactosylceramide:60% brian PS:0.5% Cy5-PE (Avanti, 810345).

All lipids were purchased from Avanti Polar Lipids as described in the Reagents and Antibodies section. Lipid mixtures were dissolved in chloroform in glass vials. Chloroform was evaporated under a stream of nitrogen gas to produce a lipid film on the glass surface, followed by further drying in a vacuum oven for 1 hour. Dried lipid films were rehydrated in liposome buffer containing 20 mM HEPES (pH 7.4), 100 mM KCl, 0.5 mM TCEP at a final concentration of 1 mg/ml (∼1.2 mM). For PS mixtures, liposomes were formed by three freeze (liquid N2)–thaw (37°C water bath) cycles. Lipid aggregates were removed by a brief centrifugation (500xg for 5min). In the case of GC/PS mixtures, lipid nanotubes were formed by a brief vortexing instead of freeze-thaw cycles, as descried (44, 45). All the liposomes, including nanotubes, were stored in the dark at 4 °C to avoid photooxidation and used within one week. The detailed protocol was deposited in protocols.io (DOI: dx.doi.org/10.17504/protocols.io.6qpvr612ovmk/v1)

### Liposome binding

#### Co-sedimentation analysis

LRRK2-FL or RCKW (300nM) were incubated in Beckman microfuge tubes at 37^0^C with PS liposomes (20 μM) in the absence or presence of different molecules as indicated in the main text for 30 minutes. The mixtures were then spun at 49,000 rpm (100,000 × g) for 20 min in a Beckman Optima TLX ultracentrifuge. Pellets were resuspended with the same volume of protein buffer as the supernatant, and analyzed by SDS-PAGE and Coomassie Blue staining.

#### Confocal fluorescence microscopy analysis

For figure 1, Rhod-PE labeled PS liposomes (20 μM) were incubated with different concentrations of LRRK2 as indicated. For figure 4, Cy5-PE labeled GC/PS nanotubes (20 μM) were incubated with GFP-LRRK2 (100 nM) in 35-mm glass bottom dishes (MatTek Corp) at 37^0^C for 30 minutes. Images were captured with a Spinning disk confocal (SDC) microscopy at room temperature on a Nikon Ti-E inverted microscope using the Improvision UltraVIEW VoX system (Perkin-Elmer). Excitation wave lengths used were 561-nm (Rhodamine), 640-nm (Cy5) and 488-nm (GFP). All images were analyzed with ImageJ. The detailed protocol was deposited in protocols.io (DOI: dx.doi.org/10.17504/protocols.io.yxmvmndqng3p/v1)

### Liposome tubulation

#### Confocal fluorescence microscopy analysis

WT and mutant LRRK2-FL or RCKW were used at the concentration of 300 nM unless otherwise indicated in the text. Liposome concentration was 20 μM. LRRK2-liposome mixtures in the buffer used for the dialysis of the purified proteins (20mM HEPES 7.4, 150 mM NaCl, 2.5 mM MgCl2, 5% Glycerol, 2mM DTT, 20 μM GDP) were prepared in a PCR tube and immediately deposited (6-10μL) on 35-mm glass bottom dishes in the absence or presence of different nucleotides as indicated in the main text and figure legends. After 30 minutes incubation at 37^0^C, images were captured with a SDC microscopy as described above. Movies were collected from time zero.

#### Quantitative analysis of liposome tubulation: differentiation and statistics of tubular and vesicular liposome structures

The Trainable Weka Segmentation plugin in FIJI was used to train a segmentation model with a representative dataset to differentiate the background, the tubular, and the vesicular structures in the Rhodamine-liposome images. The trained model could be repeatedly applied to all Rhodamine-liposome datasets using an ImageJ macro script. After a segmentation map was obtained for each dataset, a MATLAB script using area and shape criteria was applied to identify the objects that were incorrectly assigned as tubules by the Weka plugin and reassign them as vesicles. The same script then proceeded to calculate the ratios of the total area of the tubules over the sum of such area and the total area the vesicles. These results were tabulated for all categories of datasets and from them statistics were calculated for each category. The trained Weka model, the ImageJ script, and the MATLAB script are all available for download at: https://github.com/linshaova/tubule-v-vesicle.git

#### Negative stained electron microscopy (EM) analysis

LRRK2-induced liposome tubulation was performed directly on carbon-coated grids. Briefly, a glow discharged EM grid was placed into a 35-mm glass bottom dish. LRRK2 (300nM)-liposome (80μM) mixtures were prepared in a PCR tube (see above for the confocal microscopy analysis), immediately applied (6μL) to the grid and incubated at 37^0^C for 30 minutes, followed by 2% uranyl acetate staining. The grid was then blotted on filter paper and dried. Images were collected using a Talos L 120C TEM microscope at 80 kV with Velox software and a 4k × 4K Ceta CMOS Camera (Thermo Fisher Scientific).

The detailed protocol was deposited in protocols.io (DOI: dx.doi.org/10.17504/protocols.io.e6nvwkx2dvmk/v1)

### Fluorescence recovery after photobleaching (FRAP)

For FRAP experiments, SDC microscopy was used as described above. Bleaching was performed using a 488 nm laser at maximum intensity. Recordings were carried out at 1 Hz, starting with a 3-picture prebleach sequence, followed by a bleach event for 500 ms, and a post-bleach sequence up to 600 s. Each FRAP experiment was performed in at least 3 ROIs from 3 independent replications. Intensity recovery traces obtained from the regions of interest were background corrected and normalized. All the data were analyzed using Prism 9, Graphpad. The detailed protocol was deposited in protocols.io (DOI: dx.doi.org/10.17504/protocols.io.5qpvobj29l4o/v1)

### Electron microscopy (EM) analysis of LRRK2-Nanotube assembles

#### Negative staining EM

LRRK2 and nanotube mixtures were prepared in a PCR tube with a total volume of 10μL with 300nM LRRK2 and 20μM lipid nanotubes. After incubation at 37^0^C for 30 minutes, 6μL of the sample was applied to a discharged grid and adsorbed on the grid for 5 min at room temperature. The EM grids were negatively stained with 2% uranyl acetate as described above. Images were collected using a Talos L 120C TEM microscope as above.

#### Cryo-EM

Freshly purified LRRK2 was dialyzed into a low salt buffer (20mM HEPES 7.4, 90 mM NaCl, 2.5 mM MgCl2, 7% Glycerol, 2mMDTT, 20 μM GDP). After dialysis, LRRK2 (2 μM) was first incubated with MLi-2 (5 μM) for 10 min on ice, then added to lipid nanotubes (20 μM lipids) and further incubated for 1 hour at room temperature in the additional presence of 1mM GTP (Total volume of the mixture was 12 μL). LRRK2 samples (4µl each) were then applied to C-flat™ holey carbon gold grids (CF-1.2/1.3-3Au) which had been previously glow-discharged (15 mA for 45s) with the PELCO easiGlow™ Glow Discharge Cleaning System. Sample-loaded grids were plunge-frozen in liquid ethane-propane mixture using a Vitrobot Mark IV (FEI) with the following parameters: blot force, 0; blot time, 1 s; wait time, 30 s; drain time, 0 s; humidity, 100%. CryoEM micrographs were collected on a Titan Krios transmission electron microscope (Thermo Fisher Scientific) operating at 300 kV, equipped with a post column GIF Quantum energy filter and a Gatan K3 Summit DED camera (Gatan, Pleasanton, CA, USA). Data collection was performed with the SerialEM software (46). Movies were recorded in super-resolution mode with a physical pixel size of 1.098 Å (super-resolution pixel size is 0.549 Å) and a defocus range of -1 to -3 µm. The total dose of ∼60.6 e− Å−2 was attained by using a dose rate of ∼23.5 e− pixel−1 s−1 across 43 frames for 2.58 s total exposure time. The initial drift and beam-induced motions was corrected using MotionCor2 (47).

The detailed protocol was deposited in protocols.io (DOI: dx.doi.org/10.17504/protocols.io.3byl4bmdzvo5/v1)

### Generation of membrane tubules pulled from giant unilamellar vesicles (GUVs)

#### Preparation of GUVs

GUVs were prepared following the lipid-covered silica bead method described previously (48). Briefly, dioleoyil-phosphatidylcholine (DOPC), dioleoyil-phosphatidylserine (DOPS) and dioleoyl-phosphoethanolamine labeled with Atto 647N (Atto 647N DOPE) at 59.9:40:0.1 mol% were dissolved in chloroform at a final lipid concentration of 0.5 g/L. The lipid mixture was then dried in vacuum for at least two hours in an amber glass vial to form a dried lipid film, followed by hydration in 25 mM HEPES pH 7.4 buffer to form a suspension of multilamellar vesicles (MLVs) at 0.5 g/L. 20 mL of the MLVs solution was mixed with 2 mL of 40 mm silica beads (Microspheres-Nanospheres, USA), deposited on parafilm and dried in vacuum for at least one hour. The dried films over the silica beads were then initially hydrated in a 1 M trehalose solution for 15 minutes at 60 °C in a home-made humidity chamber and deposited in the observation chamber (48). Finally, the observation chamber was gently stirred manually for 1 minute to promote the detachment of the hydrated GUVs from the support silica beads.

#### Pulling membrane tubules from GUVs

Closed glass micropipettes prepared with a P-1000 micropipette puller (Sutter Instruments, USA) were used to pull lipid membrane tubules from the GUVs by direct contact between the tip of the micropipette with the GUVs. To move the micropipettes, the XY position was controlled inside the microscopy chamber using a micro-positioning system (MP-285, Sutter Instrument, Novato, CA, USA). GFP-LRRK2 was added to the observation chamber before tube pulling at the concentrations indicated in each panel, with or without different nucleotides (5 mM). The detailed protocol was deposited in protocols.io (DOI: dx.doi.org/10.17504/protocols.io.j8nlkw7p5l5r/v1)

### Quantification of the GFP-LRRK2 sorting ratio

The sorting ratio is defined by the relative change of the membrane area occupied by one GFP-LRRK2 molecule. Sorting ratios dependent on membrane curvature were calculated by the ratio between the GFP-LRRK2 and the Atto 647N DOPE fluorescence on the surface of the tubule and on the GUV with the following equation:

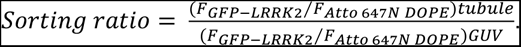

*F_GFP−LRRL2_⁄F_Atto 647N DOPE_* was measured as the ratio of the integrated fluorescence from the fluorescence plot profiles of the protein and membrane signals, respectively, neglecting the polarization factor (49).

### Quantification of the tubule radius

The radius of the lipid tubule was calculated from a calibration method based on fluorescence intensity of a flat lipid film deposited on the glass coverslip (49). The flat lipid film was used to calculate the density of the membrane fluorescence signal *π_0_.* The radius of the tubules was then calculated from the total fluorescence per unit length of the tubules *F_l_* as follows *r* = 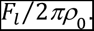

### Generation of membrane tubules by lipid-covered silica beads rolling

Lipid tubules were produced using the methodology described (50) with small modifications. Briefly, a microfluidic Ibidi sticky-Slide VI 0.4 device was positioned on top of a coverslip previously passivated with bovine serum albumin (BSA), and the lipid-covered silica beads were added to the inlet of the microfluidic chamber without the pre-hydration step. After beads addition, the microfluidic chamber was gently tilted with the outlet to the bottom, letting the beads moved in the microfluidic channel from inlet to outlet forming the tubules. The detailed protocol was deposited in protocols.io (DOI: dx.doi.org/10.17504/protocols.io.n92ldpoe7l5b/v1)

## Acknowledgments

We thank Shawn Ferguson for discussion, suggestions and critical reading of the manuscript, Karin Reinisch for the kind gift of the human LRRK2 clone, Andres Guillen-Samander, Michale Hanna and other member of the De Camilli lab for discussion, Frank Wilson and Alina Vulpe for technical assistance. We acknowledge the support of the cryo-EM facilities of the Yale West Campus and of the EM facility of the Center for Cellular and Molecular Imaging of the Yale School of Medicine. This work was supported in part by grants to P.D.C from the NIH (DA018343), the Kavli Institute for Neuroscience, the Parkinson’s Foundation, and the Aligning Science Across Parkinson’s grant ASAP-000580 through the Michael J. Fox Foundation for Parkinson’s Research (MJFF) and grants from the Swiss National Fund for research (FNS) # CRSII5_189996, #310030_200793 and European Research Council Synergy grant #951324-R2-TENSION to AR. For the purpose of open access, the authors have applied a CC BY public copyright license to all Author Accepted Manuscripts arising from this submission. S.C was supported by a fellowship from the Parkinson’s Foundation. P. De Camilli serves on the scientific advisory board of Casma Therapeutics. JE acknowledges an EMBO-Long Term Fellowship (ALT 989-2022).

## Supporting Information

**Fig. S1.**
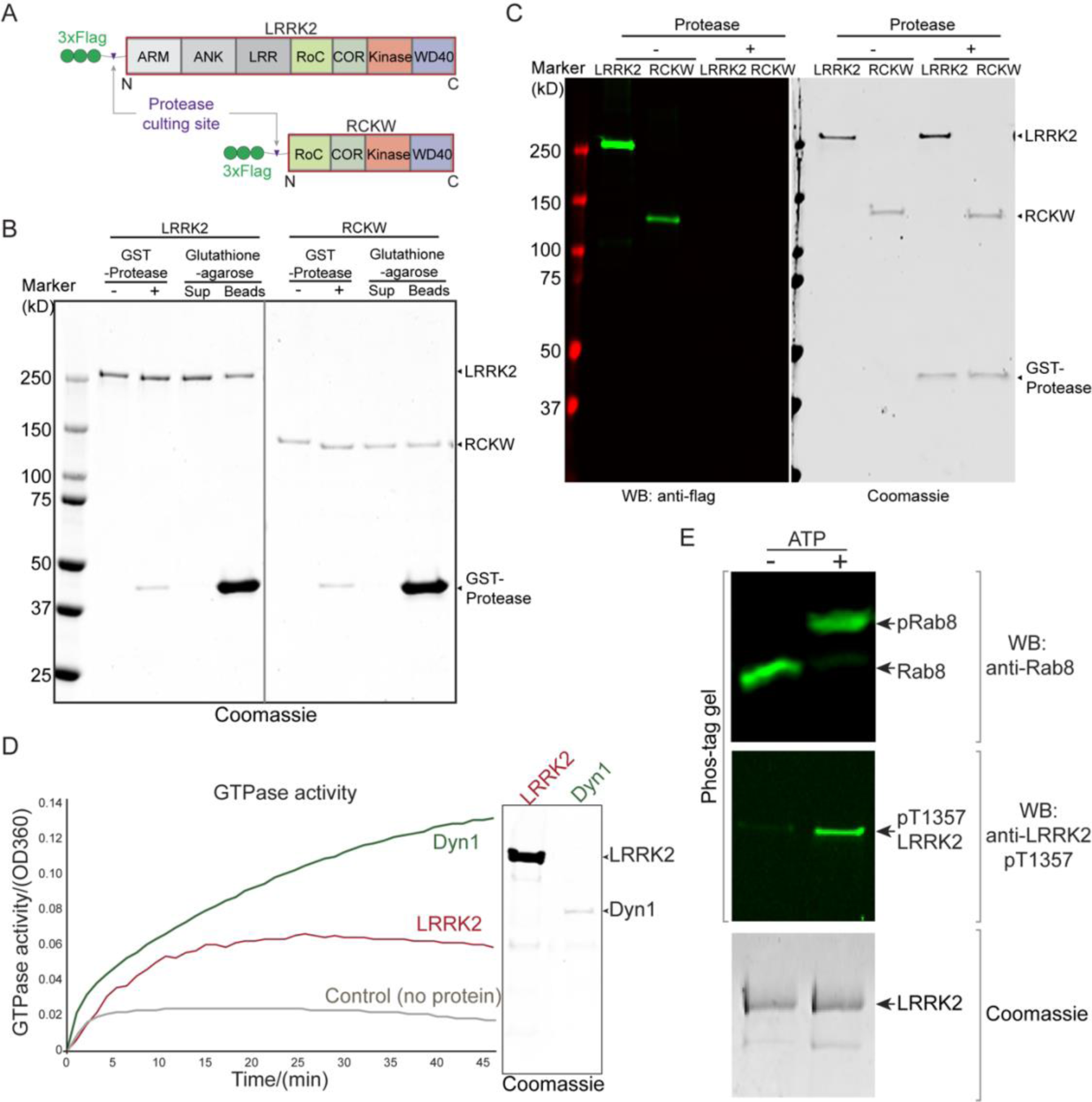
Purified LRRK2 is enzymatically active. (A) Domain cartoon of the purified proteins used in the functional assays. (B) Coomassie-stained SDS gels showing fractions obtained during the purification of recombinant full-length LRRK2 and RCKW fragment from Expi293 cells. Proteins were purified on anti-flag M2 resin, eluted from the beads using the FLAG peptide, followed by GST-Protease to remove the tag and further purified by removing the GST-protease by Glutathione-agarose. LRRK2- and RCKW-containing supernatants (Sup) were dialyzed before use. (C) Anti-FLAG tag Western blot (WB) and Coomassie-stained gel showing successful removal of the flag tag. (D) *In vitro* GTPase activity assay. The GTPase activity of LRRK2 (9 µM) and dynamin 1 (Dyn 1) (0.8 µM) were measured by phosphate release at saturating concentrations of GTP (0.5 mM). Left, LRRK2 has detectable GTPase activity but a much lower activity than dynamin 1. Right, Coomassie-stained SDS-PAGE showing the proteins used at the same molar ratio used for the GTPase assay. (E) Protein kinase activity assay. LRRK2-mediated phosphorylation of recombinant Rab8 (upper), and autophosphorylation of LRRK2 (middle), were analyzed by Phos-tag gels using an anti-Rab8 or an anti-LRRK2 phospho-specific (pT1357) antibody, respectively. Coomassie Blue stained gels of LRRK2 samples used for the assay are shown at the bottom of the figure.

**Fig. S2.**
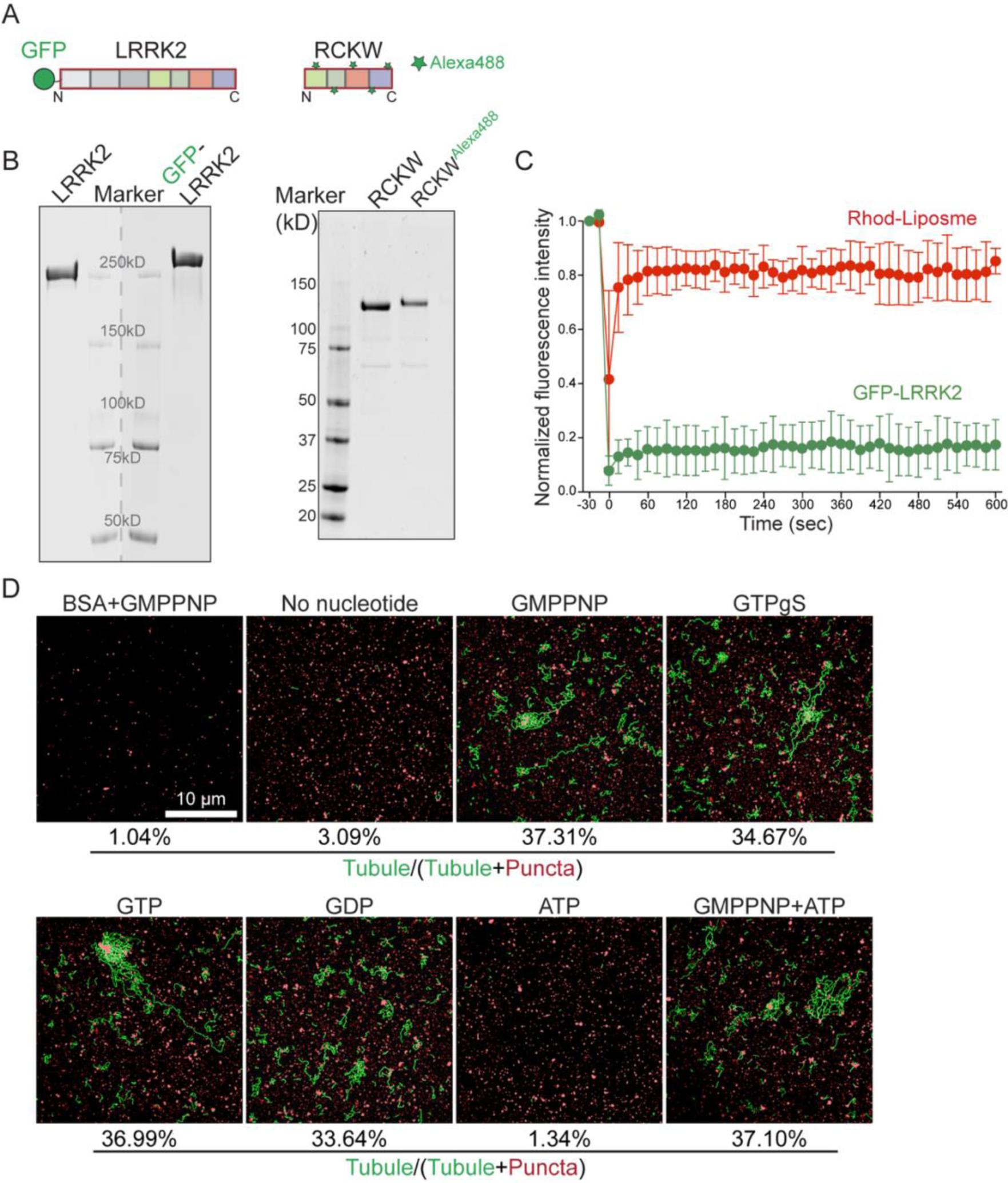
Characterization of LRRK2-induced liposome tubulation. (A) Proteins used in the tubulation assays. (B) Coomassie Blue-stained SDS-gels showing tagged proteins used for the assays and, as a control of motility, the corresponding untagged proteins. (C) Plot of the average fluorescence intensities after photobleaching of multiple ∼2mm segments on different tubules. Normalized fluorescence intensities are plotted versus time (s). Values represent mean ± S.D (n=3). (D) Representative processed images highlighting LRRK2-induced liposome tubulation in the absence or presence of different nucleotides. Values below the images represent the percent area of total liposome fluorescence accounted for by the tubules in the micrograph shown.

**Fig. S3.**
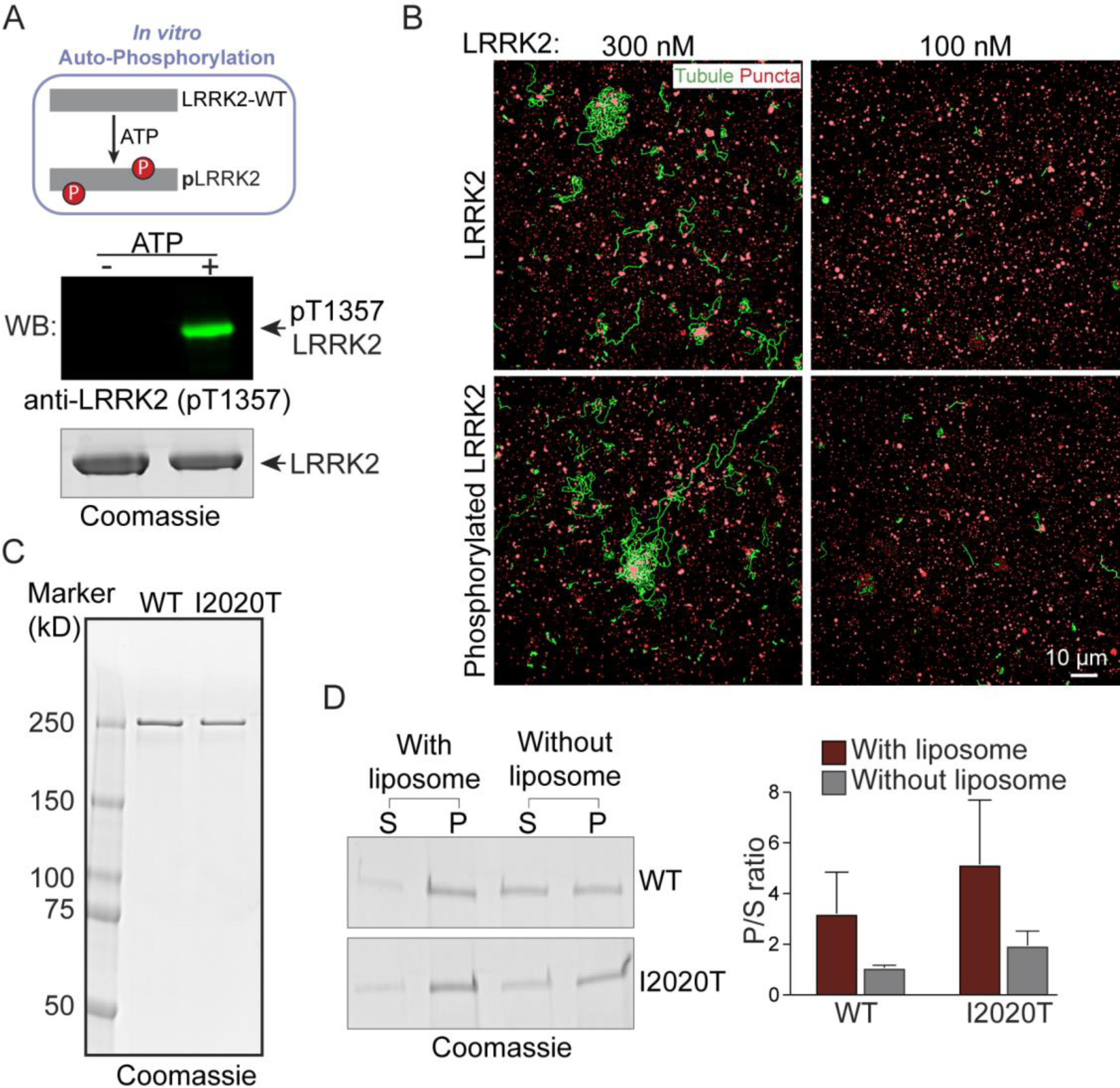
*In vitro* LRRK2 autophosphorylation and liposome binding of mutant LRRK2. (A) *In vitro* LRRK2 autophosphorylation assay. LRRK2 autophosphorylation was initiated by adding 1mM ATP to the phosphorylation mixture and validated by Western blotting using a LRRK2 phospho-specific (pT1357) antibody. A Coomassie blue stained gel of the LRRK2 proteins used is shown at the bottom. (B) Processed images of liposome tubules induced by LRRK2 (top panels) or autophosphorylated LRRK2 (bottom panels). (C) Coomassie blue-stained gel showing purified wild type (WT) and mutant LRRK2 (I2020T) proteins. (D) Left, Coomassie blue-stained gel of pellets (P) and supernatants (S) of liposomes incubated in the presence or absence of full length wild-type (WT) or mutant (I2020T) LRRK2 and then subjected to centrifugation. Right, Quantification of the intensities of gel bands in pellets and supernatants (n=3). Bars represents ratios between proteins present in the pellets and in the supernatant. Values represent mean ± S.D (n=3).

**Fig. S4.**
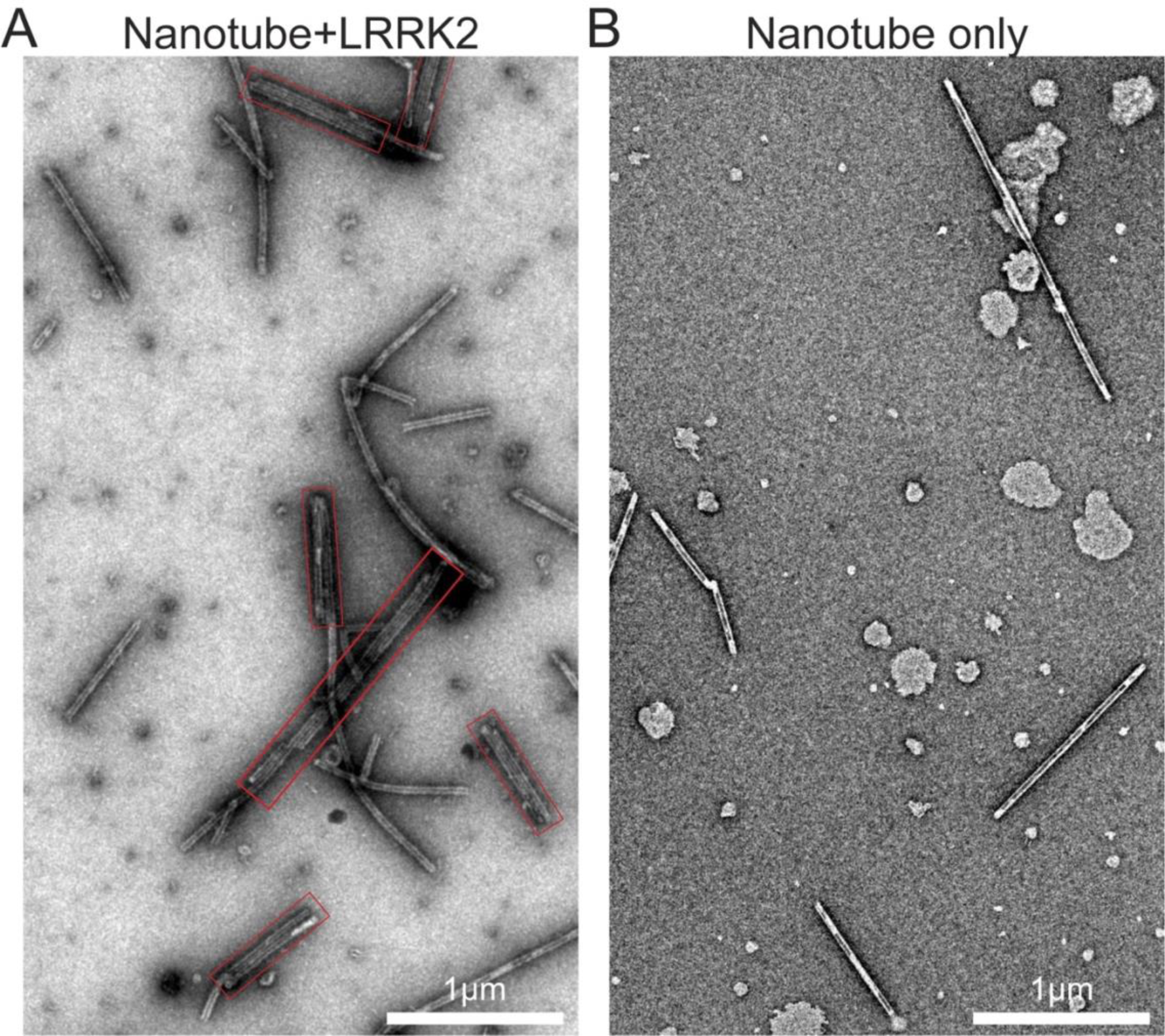
LRRK2 binds to lipid nanotubes. (A) Low magnification EM images of negatively stained nanotubes incubated in the presence of LRRK2. Note presence of nanotube bundles induced by LRRK2 (red boxes). (B) Nanotube only control.

**Movie S1 (separate file).** Confocal time-lapse imaging showing Rhod-labeled liposome tubulation induced by GFP-LRRK2. Images were collected at a rate of 1 frame every 20 seconds. Scale bars, 10 µm.

